# Transcriptomic changes in the pituitary of female mice with androgen excess through dihydrotestosterone (DHT) treatment

**DOI:** 10.1101/2021.12.31.474680

**Authors:** Serene Joseph, Sara Divall, Sheng Wu

## Abstract

Androgen excess in women is associated with the development of PCOS and its abnormalities. The Hypothalamus Pituitary Ovarian axis signaling is altered with excessed androgens, leading to anovulation and infertility. Previous studies in the lab have shown that AR signaling in the pituitary alters gonadotrophin release. Hence, the present pioneering study was an approach to determine the transcriptomic changes responsible to the phenotype seen with DHT excess. RNA seq data showed that 583 genes were differentially regulated by DHT in pituitary, of which 344 were upregulated and 239 downregulated. Pathways involved for these genes included endoplasmic reticulum, Golgi apparatus, calcium signaling and vesicles. Meanwhile, Transcriptional factor analysis showed that majority of the genes changed had Androgen responsive elements.

## Introduction

Intact androgen receptor signaling in various tissues of the reproductive axis is important for normal reproductive function and fertility. Androgen excess is also associated with reproductive and metabolic dysfunction in female humans and mice. Women with hyperandrogenemia experience associated hyperinsulinemia and polycystic ovarian syndrome (PCOS), characterized by menstrual irregularities, anovulation, hirsutism, and impaired glucose tolerance/diabetes mellitus (*2*). Lean female mice given low dose dihydrotestosterone (DHT)– a model of lean PCOS--also exhibit reproductive and metabolic manifestations including hyperinsulinemia, altered insulin signaling in metabolic and reproductive tissues, and infertility (*3*).

Female mice with complete knockout of the AR receptor are phenotypically normal, but have reduced fertility and litter size (*4–6*). Knockout of AR in gonadotrophs indicate that AR is important for the maintenance of normal levels of gonadotrophins to promote proper recruitment of follicles and in ovulation (*7*). Androgen receptors (AR) play a pivotal role in the pathogenesis of PCOS (*8*), and are expressed in PCOS-affected organs such as gonadotrophs, liver and ovary (*7, 9*). Gonadotroph specific ARKO female mice treated with DHT exhibit improved fertility and regular estrous cycling compared to wild type female mice treated with DHT, suggesting that excessive AR signaling modifies gonadotropin release(*1*) Thus, excessive or defective AR signaling affects gonadotropin synthesis and release. Thus, to understand the pathways affected by excess dihydrotestosterone (DHT) in the pituitary in an unbiased manner, RNA sequencing was done in pituitary glands collected from female wild type mice and mice treated with DHT. Comparison of the transcriptomes of these groups allow us to isolate components of the transcriptome that are directly affected by androgen receptor signaling in the gonadotroph.

## Results and Discussion

### WT and WTDHT samples were distinctly separated by DEGs

RNA Sequencing data comparison was done to determine the effect of DHT in the presence of AR receptors in the pituitary. The treatment involved the administration of DHT, and ORA analysis of Gene ontology included biological processes, Cellular component and Molecular function.

583 genes (DEGs) were differentially expressed in the pituitaries of the female WTDHT mice compared to the WT mice, out of which 344 genes were upregulated (positive log FC) and 239 downregulated (negative log FC) (Figure 1). Genes whose threshold were true are in blue and false in red.

**Figure 1:**
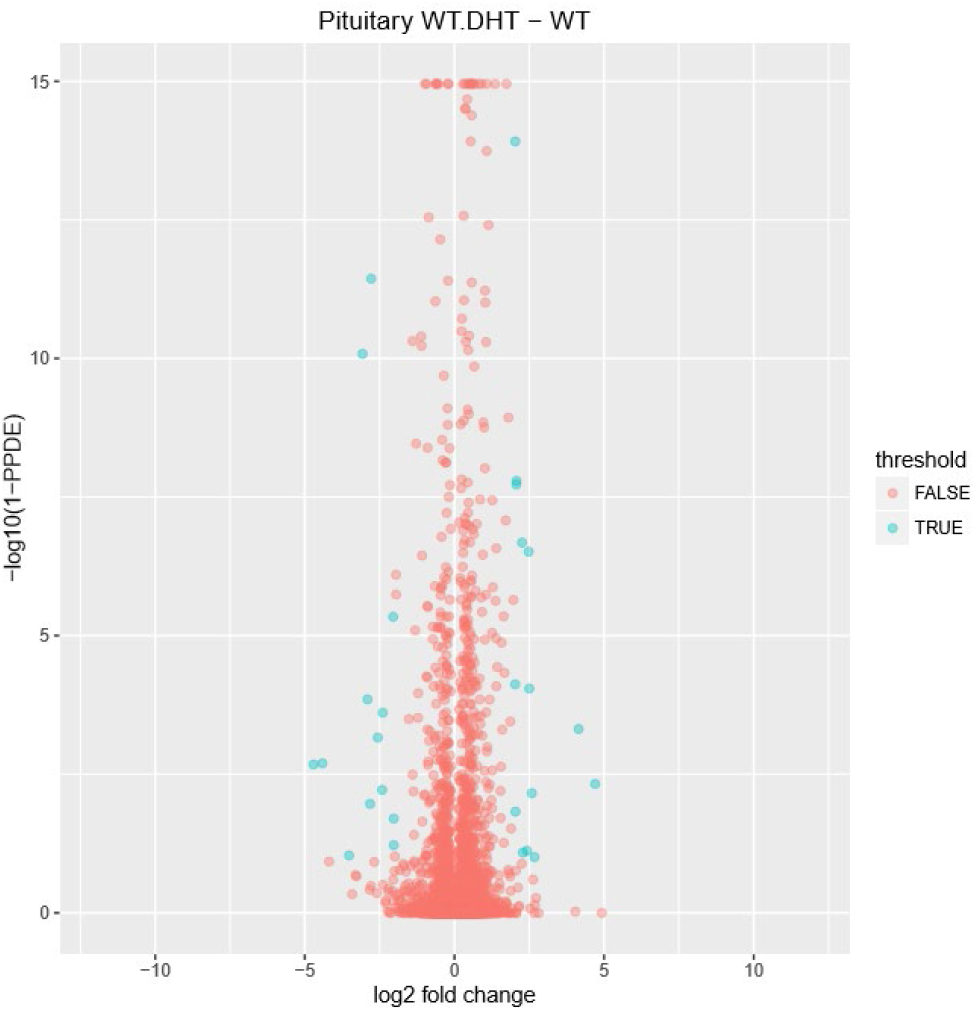
Volcano plot showing the gene expression in pituitary WTDHT compared to WT. 583 genes were differentially regulated (DEG) in the pituitaries of the female WTDHT mice compared to the control (WT mice), out of which 344 genes were upregulated (positive log FC) and 239 downregulated (negative log FC),

583 DEGs identified from Figure 1 were used to generate heatmap with mean subtracted signals. As shown in Figure 2, these DEGs could well-separate WT and WT DHT samples, confirming their potential to probe androgen-related signaling pathways in pituitary.

**Figure 2:**
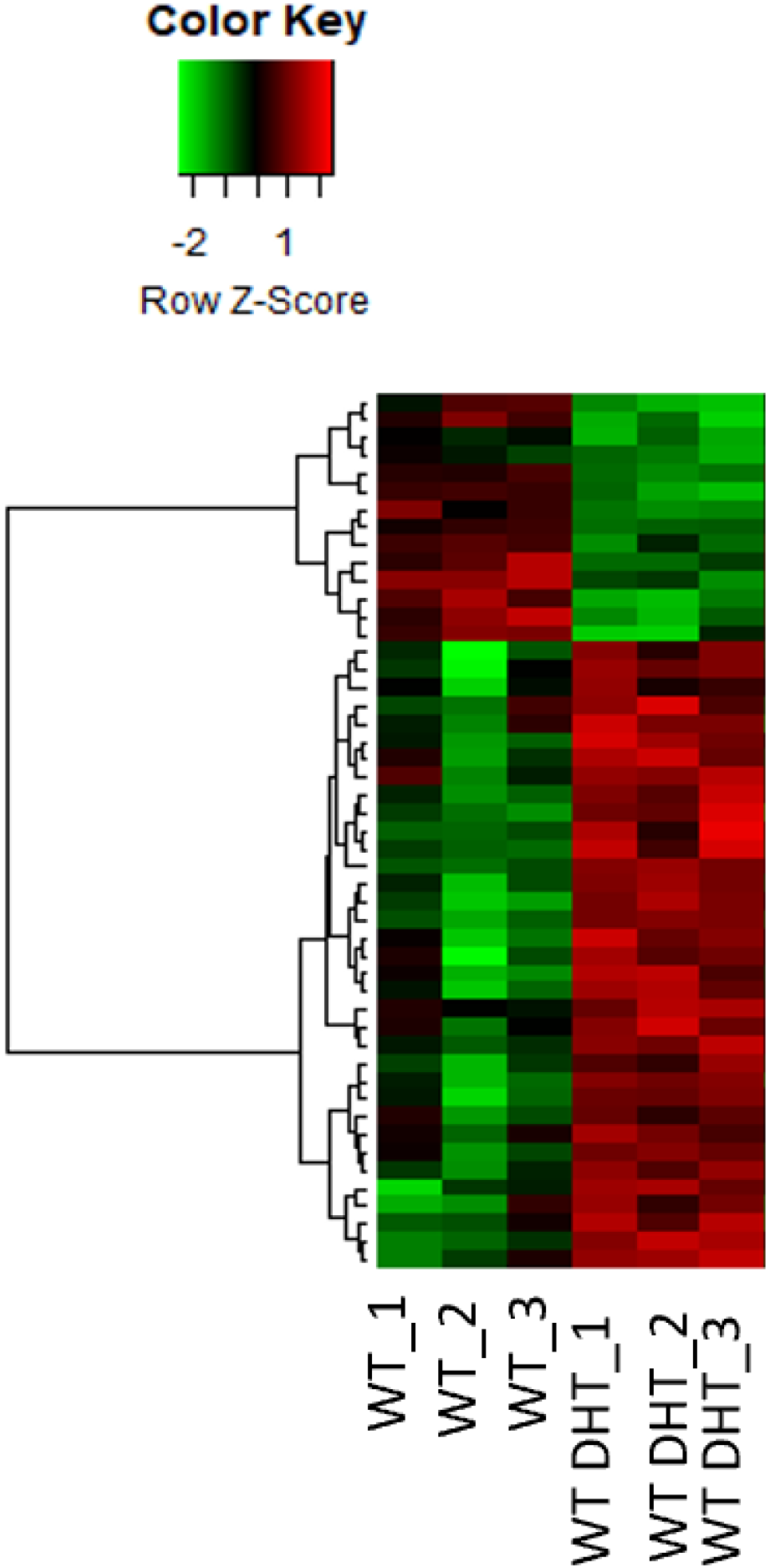
Heat map of genes differentially regulated by DHT in the pituitary.

### Pathways overrepresented by DHT

ORA analysis of the DEG was done on Webgestalt 2019 and the biological processes, molecular function and Cellular components that were overrepresented are summarized here.

Biological processes that were overrepresented in the pituitary after DHT treatment were related to protein folding, cell adhesion, negative regulation of organelle localization, epithelial cell migration, anterograde trans-synaptic signaling, peptide transport, establishment of mitotic spindle orientation and regulation of exit from mitosis. The most significantly upregulated biological processes in the order of their enrichment are microtubule anchoring at the centrosome, mitotic cell cycle arrest, cell substrate junction assembly, maintenance of location in cell, skeletal system morphogenesis, epithelial cell migration and kidney development. The downregulated processes are chaperone mediated protein folding, protein binding, response to unfolded protein, organic substance catabolic process.

The molecular functions that were overrepresented upon treatment with DHT in the pituitary were related to transporter activity, which narrowed down to organic acid: sodium symporter activity, Factors that regulate molecular function were also overrepresented, and specifically narrows to GTPase activator activity. Other overrepresented functions were related to amyloid beta binding, protein dimerization activity, ATP dependent protein binding, glutamate receptor binding, transcription factor binding, identical protein binding, heat shock protein binding, calcium binding, adenyl deoxyribonucleotide binding, chaperone binding. The downregulated functions were related to ATP dependent protein binding, unfolded protein binding, chaperone binding, heat shock protein binding, isomerase binding, ubiquitin protein ligase binding, purine nucleotide binding and drug binding. The upregulated functions were related to tubulin binding, protein kinase binding, calcium ion binding, zinc ion binding, protein protein complex binding, transporter binding, cytoskeletal protein binding, signaling receptor binding and molecular function regulator.

Cellular components that were most enriched upon DHT treatment narrowed to extracellular exosome and kinocilium. Besides this component such as inotropic glutamate receptors were highly enriched. Overrepresented pathways related to the upregulated genes are pericentriolar material, extracellular exosome, inotropic glutamate receptor complex, integral component of the presynaptic membrane, inotropic glutamate receptor complex. The downregulated pathways were related to tubular endosome, chaperone complex, integral component of Golgi membrane. The overall overrepresented cellular components with their genes, enrichment ratio and FDR values are given in Table 1.

### Pathways related to gonadotrophin secretion and their interaction networks

To get a mechanistic insight into the pathways by which excess DHT alters the pituitary and gonadotroph secretion, relevant pathways to gonadotrophin secretion were analyzed in more detail. With relevance to gonadotroph secretion, the altered cellular components recognized by WEBGESTALT 2019 were related to Gene Ontology terms like cytoplasmic vesicle, endosomes, adherens junction and integral component of the endoplasmic reticulum and the Golgi apparatus.To further explore the calcium and pituitary signaling activities affected by androgen, we plotted the string network of genes related to calcium ion binding (Figure 3a), and genes in organelle sub compartment (Figure 3b). As most of these genes have little been known to be related to androgen so far, this study rolls out the potential biomarkers which are related to excess androgen in pituitary.

**Figure 3a:**
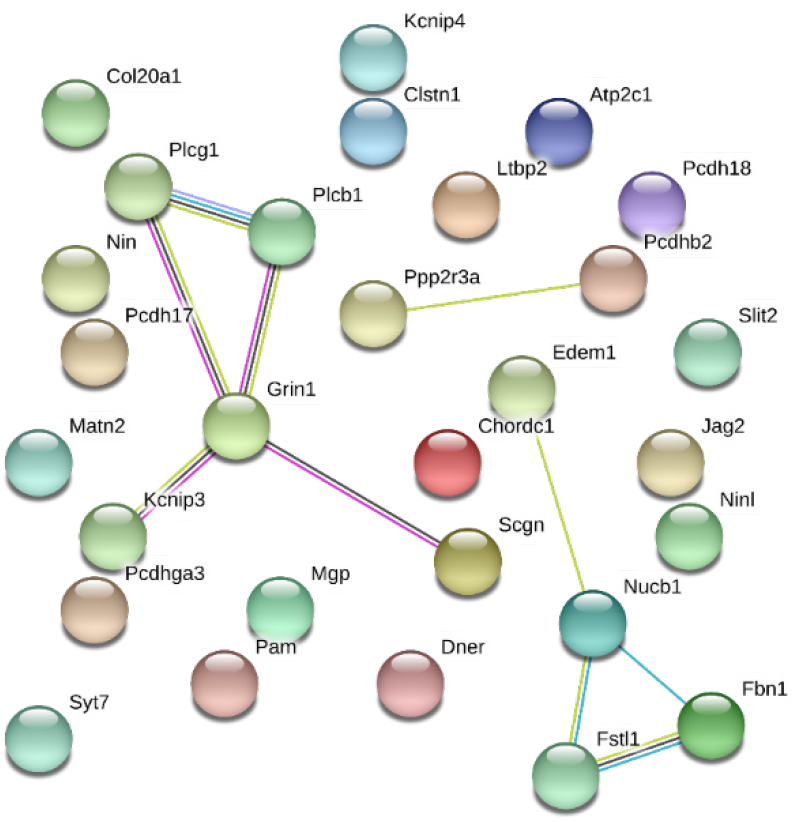
String network showing interactions between genes that are related to calcium ion binding.

**Figure 3b:**
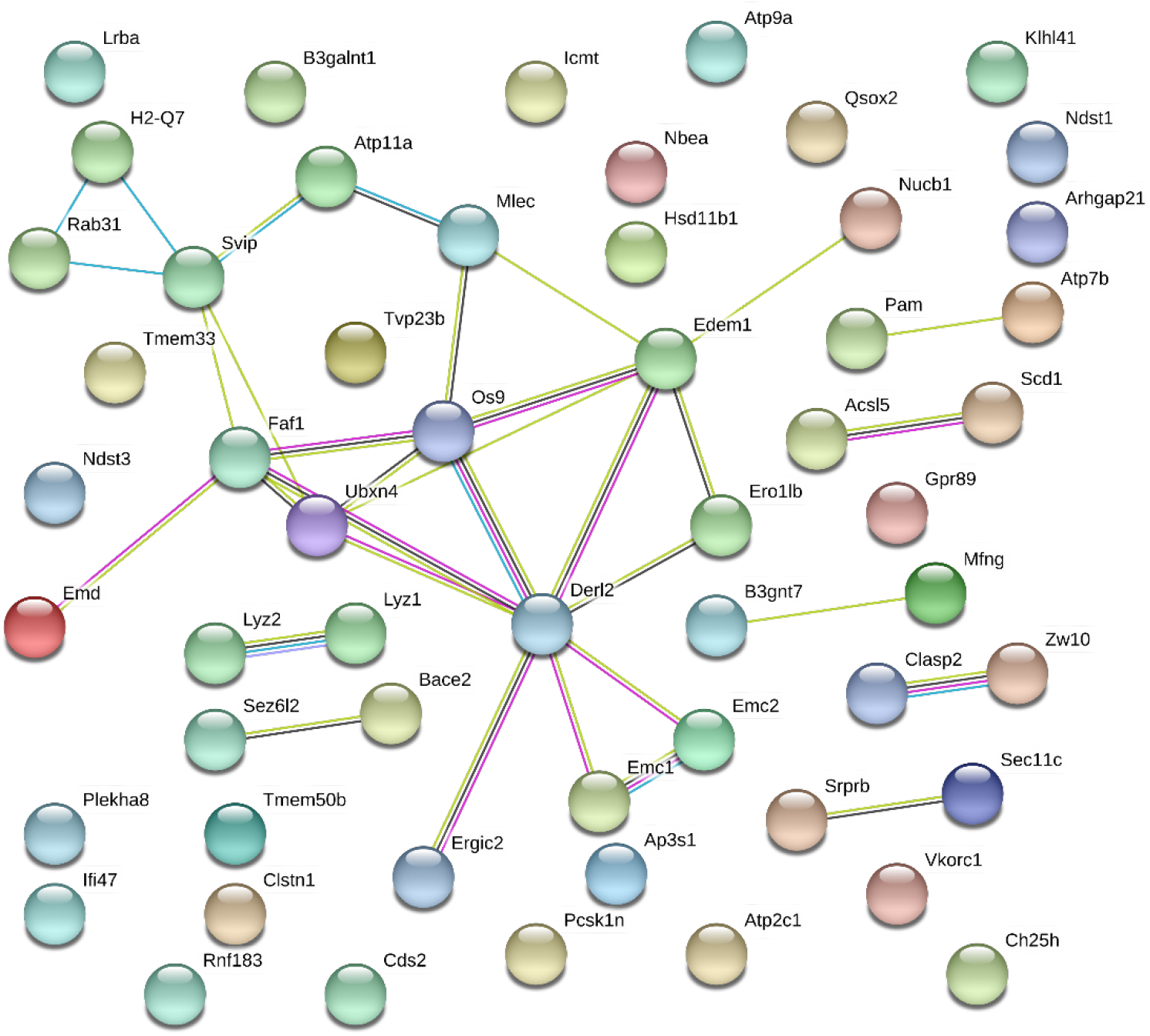
String network showing interactions between genes involved in organelle sub compartment.

#### Transcription Factor Analysis

The genes DEG between WTDHT/ WT were compared to genes having full AR TF and half AR TF sites as determined by CHIP -sequencing identified (*10*). It was seen that most genes differentially regulated by DHT treatment had both half and full ARE (androgen response element) (313 genes) or either full (34 genes) and half (98).

## Discussion

Studies indicate a potential role for AR signaling for normal gonadotropin function in the normal (*7*) and hyperandrogenic state (*1*). However, the gene networks and potential cellular pathways affected by altered AR signaling in the gonadotroph is not known. Transcriptomic analysis with determination of represented pathways in the pituitary glands from female mice subjected to chronic low dose (~2 fold) dihydrotestosterone treatment provides genomic mechanistic insight underlying the observed phenotype of attenuated LH surge upon GnRH stimulation and impaired estrous cycling (*1*). Synthesis and secretion of gonadotrophins is complex, involving transcription, translation in the endoplasmic reticulum, glycosylation and glycan remodeling, hormone packaging in the golgi apparatus and subsequent exocytosis (*11*). There were no changes in the mRNA levels of genes encoding for LH, FSH or GNRH hormone receptors, suggesting that that the concentrations of the hormones in circulation were not affected directly by transcription, but rather by genomic changes in the organelles that mediates its synthesis and secretion.

Transcriptomic changes by DHT in the pituitary were related to cell components in the secretion of gonadotrophs like endoplasmic reticulum, Golgi apparatus, lysosomes, and secretory vesicles, along with changes in Calcium ion binding which is known to have a major role in the synthesis of proteins in the gonadotrophs along with the exocytosis of hormones from the plasma membrane were different.

The endoplasmic reticulum and Golgi apparatus are important cell organelles in which pre-protein processing and synthesis of LH and FSH are carried out. Downregulation of the endoplasmic reticulum, Golgi apparatus suggest that protein synthesis and processing may be impacted. Not much is known about the impact of DHT on endoplasmic reticulum in the pituitary, but studies in the prostate gland suggest that androgens expand unfolded protein response (*12*). While there were no changes in the basal levels of LH and FSH in the DHT treated mice, there were a significant reduction in LH secretion following GnRH stimulation. Vesicles were a highly enriched cellular component, and indispensable in the formation and release of the hormones, LH and FS which control ovulation and fertility. Rab proteins formed a major cluster in the genes enriched for vesicels. Besides calcium signaling, which is also an altered pathway, controls the release of LH, from the vesicles. PLGC1 encodes the gene which catalyzes the formation of IP3 and DAG. IP3 is an important mediator of the SERCA pump, which controls free calcium levels in the cytosol ultimately needed for gonadotrophin exocytosis. (*13*).

Animals that developed PCOS like conditions on treatment, did not have a steady release of LH in response to GnRH (*3*). Transcriptomics suggest an increase in the mRNA levels of the proteins involved in Calcium binding (an overrepresented pathway) between WTDHT/ WT, thereby likely reducing the levels of free cytosolic Ca available to promote cell signaling and the fusion of secretory vesicles to the plasma membrane. The role of DHT in impaired calcium signaling in GnRH mediated LH secretion has been proven by previous studies in the lab (*1, 14*). GEM, a small GTPase, identified to be increased in the WT-DHT animals that developed PCOS, and downregulated in the pituitaries of mice lacking AR in the pituitary (*1*) is transcriptionally regulated by the excess DHT through the AR transcriptional machinery in its promoter site. Therefore, its impairment in Calcium signalling leads to altered exocytosis of hormones in the WTDHT animals.

The changes in the calcium signaling, mediated by Gem and also the transcriptomic changes related to protein processing units like the ER and Golgi apparatus, likely changes the reproductive phenotype of the DHT treated animals, by altering the secretion of LH and FSH under conditions of GnRH stimulation.

These studies indicate a potential role for AR signaling normal and hyperandrogenic condition as in PCOS (*1*) and reveal important pathways and signalling mediators that can be studied further to understand and treat PCOS.

## Methods

### Tissue collection and disease models

Two groups of female mice were used in the experimental design: Wild type mice given DHT (WT DHT), and Wild type control. The details of DHT treatment have been described previously (*3*). Briefly, female mice, roughly 2 months of age, were implanted with either 4mm DHT (WTDHT) or vehicle (WT) pellets for 3 months, replaced every month. DHT is widely used to study hyperandrogenism, because of its inability to be converted to estrogens and its binding specificity to the AR receptor (*15*). This DHT pellet implantation increased the circulating level of DHT in the adult female animal by nearly 2-fold higher than normal (*3*). At the end of the 3 months, animals were sacrificed, tissues collected and snap frozen until further analysis.

### RNA seq and pathway analysis

RNA was prepared from the whole pituitaries using the phenol chloroform method as previously described (*16*). To determine the genes and their pathways that were altered by DHT, pituitaries from WT mice implanted with DHT pellet were compared with the pituitaries from WT mice implanted with vehicle pellet, which serve as the control. DEG (Differentially Expressed Genes) from the different treatments were subjected to pathway analysis using WEBGESTALT 2019 (WEB-based Gene Set Analysis Toolkit), an open platform functional enrichment analysis web tool (*17*). The pathway analysis was performed using all statistically significantly changed genes (ppde >0.95 (1-target FDR) and then using upregulated and downregulated genes separately to better understand their functional implications as overrepresented pathways. These analyses were done using the Over representation analysis (ORA) using FDR (False Discovery Rate) set at 0.05 using the BH (Benjamini-Hochberg) adjustment. To reduce redundancy, weighed set cover criteria was used to tabulate results. The overrepresented pathways were subjected to PPI (protein protein interactions) interactions using STRING to determine interactions between the genes using a set confidence using criteria like text mining, experiments, databases, co-expression, neighborhood, gene expression and co-occurrence.

AR as a transcription factor, binds to the Androgen response element (ARE) in the promoter regions of genes and influences their transcription by either promotion or repression. With this important function of the AR receptor in mind, a transcription factor analysis was done for the DEG by excess DHT treatment in mice. Transcription factor analysis was done using WEBGESTALT 2019, with an FDR≤0.05. The DEG between WTDHT/WT were compared to genes containing full and partial ARE (tandem repeats of two hexamers) sites as identified by CHIP-sequencing (*10*).

